# Persistent Prolate Polymersomes for Enhanced Co-Delivery of Hydrophilic and Hydrophobic Drugs

**DOI:** 10.1101/796201

**Authors:** Nicholas L’Amoreaux, Aon Ali, Shoaib Iqbal, Jessica Larsen

**Author notes:** corresponding author;, 209 Earle Hall, Clemson, SC 29634.

## Abstract

Self-assembled polymersomes encapsulate, protect, and deliver hydrophobic and hydrophilic drugs. Though spherical polymersomes are effective, early studies suggest that non-spherical structures may enhance specificity of delivery and uptake due to similarity to endogenous uptake targets. Here we describe a method to obtain persistent non-spherical shapes, prolates, via osmotic pressure and the effect of prolates on uptake behavior. Polyethylene glycol-b-poly(lactic acid) polymersomes change in diameter from 175 ± 5nm to 200 ± 5nm and increase in polydispersity from 0.06 ± 0.02 to 0.122 ± 0.01 nm after addition of 50 mM salt. Transmission and scanning electron microscopy confirm changes from spheres to prolates. Prolate-like polymersomes maintain their shape in 50 mM NaCl for seven days. Nile Red and bovine serum albumin(BSA)-Fluorescein dyes are taken up in greater amounts by SH-SY5Y neural cells when encapsulated in polymersomes. Prolate polymersomes may be taken up more efficiently in neural cells than spherical polymersomes.

## Introduction

The Blood-Brain Barrier (BBB) presents one of the greatest challenges left in modern medicine. As our knowledge of therapeutic molecules and their functions increases, we must also confront how to deliver them effectively. The BBB blocks 98% of small molecules from passage, and those that do pass through must have specific properties such as hydrophobicity and are often delivered with low efficiency.^1–11^ Therapies must not only effectively deliver drugs across the BBB but simultaneously be specific, avoid immunogenic response, and minimize off target delivery.^12^ Polymeric nanoparticles, namely polymersomes, have gained traction as a potential solution due to their ability to encapsulate both hydrophilic and hydrophobic molecules, offer immune shielding, and exhibit low toxicity. Polymersomes are bilayer, membrane-bound vesicles comprised of hydrophilic and hydrophobic polymers, with a ratio of approximately 25-40% by weight hydrophilic.^13^ Based on the polymers selected, polymersomes can release due to pathophysiologic changes in the diseased area. For example, polymersomes can respond to pH,^14–21^ hypoxia,^22–24^ and temperature.^25^ Therefore, manipulation of these properties is considered a viable tool in creating drug delivery systems.

The physical properties of these nanoparticles, in our case highly monodisperse polyethylene glycol-b-poly (lactic acid) (PEGPLA) polymersomes, have been implicated as important factors in the physical mechanisms of uptake. Properties such as size, shape, charge, and material are all being studied in order to determine how to best assign these nano-vesicles with a desired uptake pathway. Natural uptake targets, such as proteins, are usually not perfectly spherical but rather exhibit a variety of morphologies. Mimicking these morphologies could allow for higher, more specific delivery that imparts less stress upon the body. For example, targeted, rod-like polystyrene nanoparticles accumulated in Caco-2 and Caco-2/HT-29 cells to a greater extent than spherical and disc like particles.^26^ Similarly, long rods of mesoporous silica covered in PEG demonstrated approximately five-fold greater uptake in HeLa cells compared to short rods and spherical PEGylated particles.^27^ The ability of HeLa cells to better internalize rod-like particles has been previously studied and reviewed.^28^ Specifically, prolate shaped nanoparticles exhibit higher aspect ratios providing greater surface area for uptake via invagination. The idea of utilizing different nanoparticle shapes for enhanced biologic uptake in a diseased area is not a new one. Kolhar, et al., found that polystyrene nanorods showed higher targeted uptake towards the endothelium compared with spherical particles, confirmed by both *in vitro* and *in vivo* assays. Furthermore, the greatest discrepancy in uptake was observed in the brains of mice, where targeted polystyrene rods were taken up in 7.5x greater amounts than targeted polystyrene spheres.^29^ We extended this application to self-assembled systems, capable of delivering more than one drug simultaneously.

Disease treatment often requires a combination, or cocktail, of drugs working synergistically in order to achieve desired outcomes. Due to the complex biochemical pathology of most diseases, treatments benefit from the ability to administer multiple therapeutics simultaneously. Specifically, diseases confined behind the BBB are only naturally susceptible to hydrophobic molecules. Many peptide-based therapies, plant-based anti-inflammatories, and antibiotics are unable to diffuse across the BBB, rendering them ineffective for treatment of brain disease. The creation of drug delivery systems, such as the one proposed in this paper, which allow the simultaneous delivery of drugs or therapeutic molecules with a range of hydrophobicity and hydrophilicity, show promise in tackling a variety of complex biological problems.

Polymersomes have been transformed to achieve a variety of shapes including rods and stomatocytes.^30,31^ More complex structures, such as nested vesicles, have also been achieved.^32,33^ The primary method of achieving these transformations is osmotic shock, creating an osmotic pressure gradient causing the movement of water molecules either into or out of self-assembled structures. Osmotic shock has resulted in shape changes through the addition of NaCl^34^, glucose^32^, or PEG^33^. Herein we report that PEGPLA polymersomes are capable of encapsulating hydrophilic and hydrophobic compounds simultaneously, and that loaded polymersomes deliver encapsulated compounds to cells more effectively than free dye alone. We confirm that subjecting PEGPLA polymersomes to 50mM NaCl solution via dialysis, as reported by Wauters, et al., results in a change in shape from spheres to prolates.^34^ Furthermore, we report that this shape change is complimented by an increase in the amount of both hydrophobic and hydrophilic compounds retained in the polymersomes.

We employed simple, yet effective methods of achieving persistent shape change in polymersomes and investigated how these changes influence the ability of the polymersomes to co-deliver drug payloads. Through the manipulation of the ionic gradient in the suspension of polymersomes, achieved via dialysis in NaCl, we show polymersomes in prolate form persist over time. Furthermore, these prolate nanoparticles were more effectively taken up by human neural cells (SH-SY5Y) in vitro. The combination of morphologically tuned prolate particles with amphiphilic encapsulation promise potential in providing minimally invasive procedures for currently untreatable diseases.

## RESULTS & DISCUSSION

The general concept for modulating spherical polymersomes to prolates by exposure to hypertonic conditions is introduced in Figure 1. Polymersomes are formed by a process of self-assembly in an aqueous solution wherein the hydrophobic portions of the diblock copolymers align to form a bilayer membrane.^13^ PEG_1000Da_-b-PLA_5000Da_ (PEGPLA) forms highly, stable spherical polymersomes via solvent injection, as described in the methods section, with properties listed in the first row of Table 1. In water, consistent sizes were observed, demonstrated by a small standard deviation in diameter and low polydispersities. Upon removal of organic solvent from the suspension of polymersomes, the polymersomes appear to approach a critical size (Figure 2) around 190 nm. *In vitro* blood-brain barrier studies indicate that nanoparticles of 200 nm or less are preferentially transported through to the abluminal side, ^37-39^ making our nanoparticles capable of similar transport behavior.

**Table 1.**
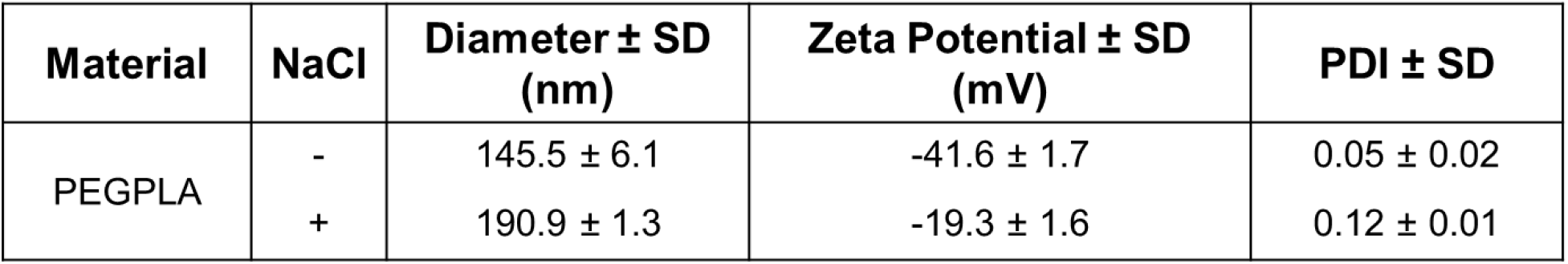
Characterization by Dynamic Light Scattering of PEGPLA polymersomes without NaCl and with NaCl at 50mM.

**Figure 1:**
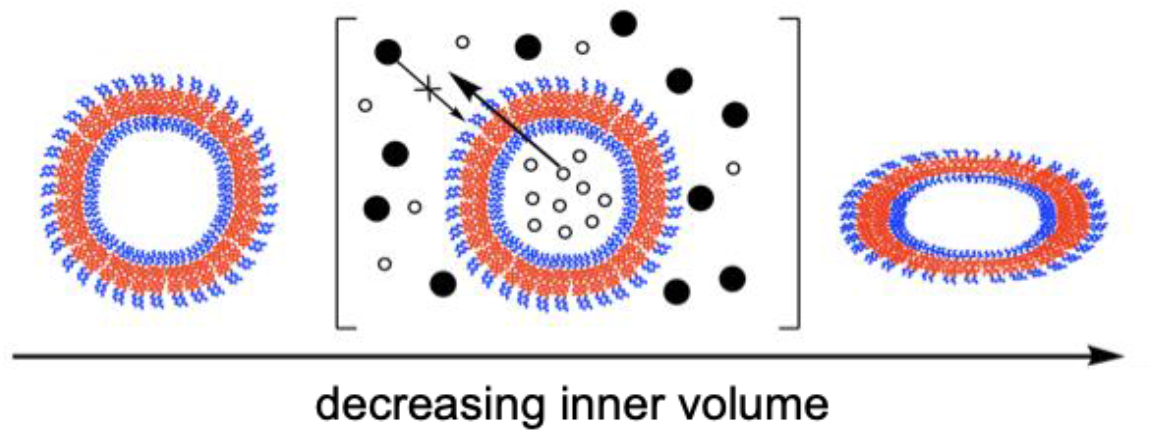
Polymersome Shape Change. Osmosis across polymersome membrane due to hypertonic conditions, resulting in a prolate shape.

**Figure 2.**
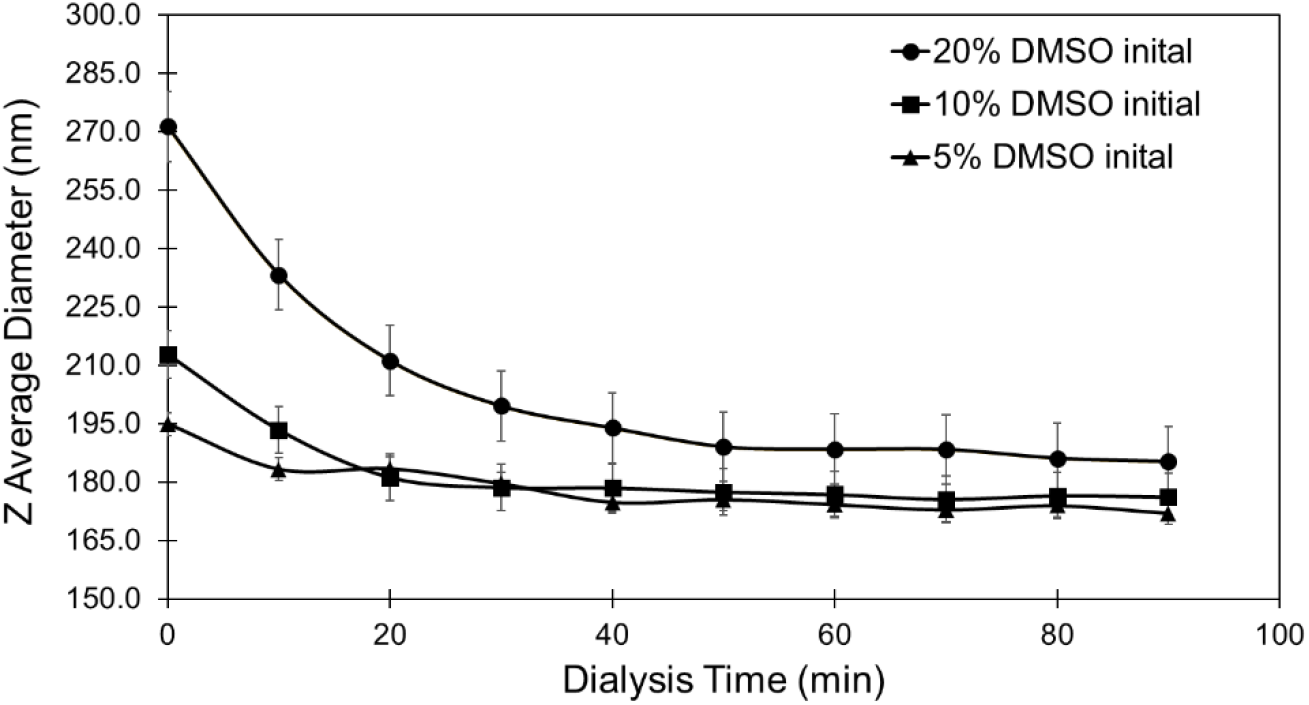
Dialysis of Spherical Polymersomes. Each sample was made from PEGPLA block co-polymer dissolved in DMSO at an equal concentration of 3mM. The 20%, 10%, and 5% samples contained3.6mg, 1.8mg, and 0.9mg of PEGPLA dissolved in 200μl, 100μl and 50μl DMSO, respectively. The samples were injected into DI water such that the final volume for each sample was 1mL. Polymersome samples were then dialyzed against DI water at dialysate volumes such that the final DMSO concentration in each sample was the same, resulting in similar final diameters.

Not unlike biological bilayer membranes, self-assembled PEGPLA polymersomes are affected by osmosis. Exposure of polymersomes to hypertonic conditions by dialysis against a salt (NaCl) solution provokes the outward diffusion of water from the core of the polymersomes^31^. In contrast to phospholipids, which make up cell membranes, the hydrophobic portion of diblock copolymers in the membrane of polymersomes are much longer, theoretically inhibiting the passage of sodium or chloride ions through the membrane to the aqueous core. The resulting effect is a decreased inner volume of polymersomes. Spherical polymersomes were dialyzed against 50 mM NaCl to create hypertonic conditions, causing a shift in thermodynamic equilibrium, resulting in a shape change. In the presence of salt, sizes became less consistent, with an increase in diameter size and polydispersity (second row, Table 1). Because dynamic light scattering assumes spherical shape^35^, an increase in perceived polydispersity could be due to an increase in aspect ratio, leading to measurements with a wider range of sizes based on the orientation of the particle in solution when the laser contacts it. Zeta potential measurements indicate that both prolates and spherical polymersomes are stable in solution, with values around or less than −20 mV.^36^

When the inner volume decreases and the surface area of the polymersomes remains constant, a shape change is observed (Figures 3, 4). Without NaCl, PEGPLA polymersomes present as spherical structures with a brush-like exterior layer, observable through the rough surface presentation in SEM. After the addition of NaCl, PEGPLA polymersomes lose their spherical morphology and appear more worm-like, indicative of a shift in shape (Figure 3). TEM images lead to the same conclusions, showing much longer and oval-like structures after the addition of NaCl. Without NaCl, PEGPLA polymersomes appear as membrane-bound spherical structures with a relatively monodisperse distribution, confirmed through dynamic light scattering (Table 1). With the addition of 50 mM NaCl, PEGPLA polymersomes become prolate structures with an aspect ratio around two, with the longest radius being approximately twice the size of the shortest radius (Figure 4). For the purpose of drug delivery applications, it is important that membrane-bound prolates, or prolate polymersomes, maintain their shape over an extended period. Prolates maintain their structure over a period of one week, as shown by minimal changes in both z-average diameter and polydispersity indices (Figure 5). Prolate polymersomes appear to have a three-day settling period, where diffusion of solvent out of the hydrophobic membrane is believed to be occurring and diameter and polydispersity index are slowly decreasing. However, after seeing a statistically significant (p< 0.05) decrease in both size and polydispersity index from day 3 to day 4, prolate polymersomes stabilize and maintain their sizes over the remaining days. It is important to note that these sizes and PDIs are still significantly larger than spherical polymersomes made without the presence of NaCl.

**Figure 3.**
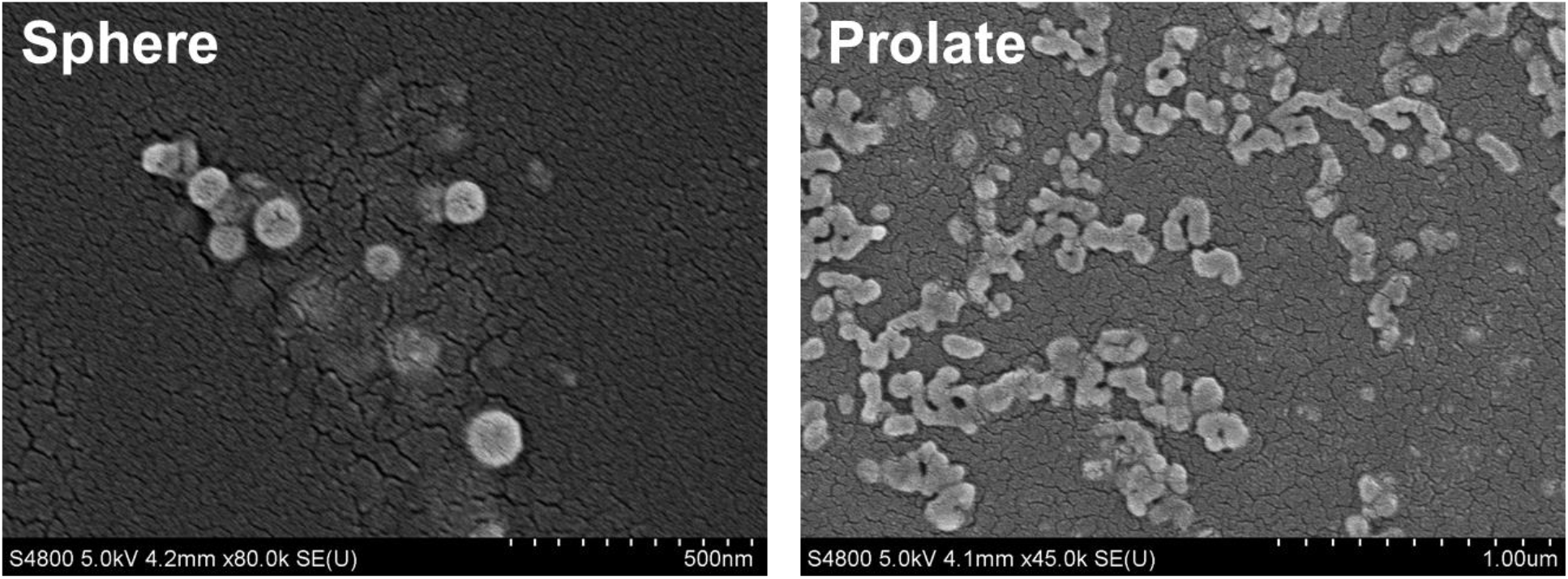
Scanning Electron Microscopy Images of Spherical and Prolate Nanoparticles. Without salt dialysis (L), SEM images display self-assembled spheres with a fuzzy core, indicative of a fully extended PEG layer. Post salt dialysis (R), SEM images display more oblong, rod-like structures(LHS Scale Bar: 500 nm; RHS Scale Bar: 1.00 μm.

**Figure 4.**
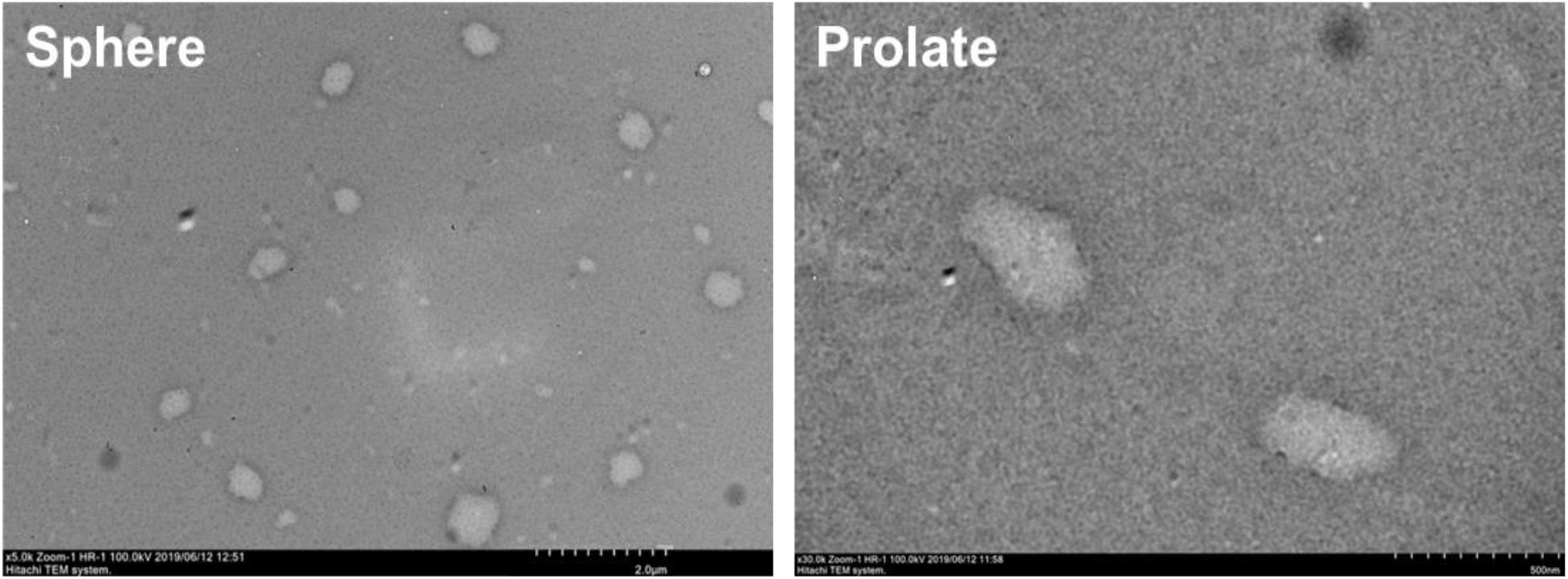
Transmission Electron Microscopy Images of Spherical and Prolate Nanoparticles. Spherical and prolate particles were dried from a 0 mM and 50 mM NaCl suspension, respectively and stained with uranyl acetate. Without NaCl, PEGPLA polymersomes appear as membrane bound spherical structures with a monodisperse distribution. With NaCl, PEGPLA are prolate structures with an aspect ratio around 2, with the longest radius being two times the size of the shortest radius. Images taken at 120kV / 60000x direct mag on Hitachi h7600 Transmission Electron Microscope.(LHS Scale Bar: 2.0 μm, RHS Scale Bar: 500 nm)

**Figure 5:**
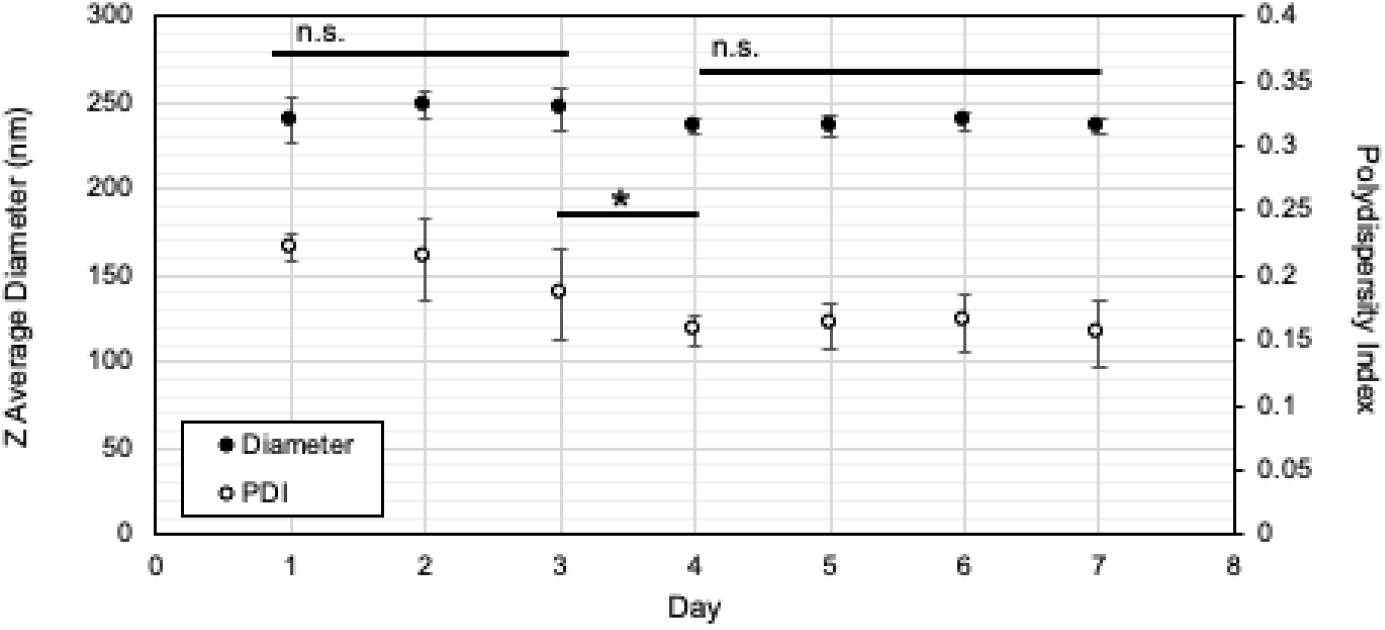
Persistent Prolate Shape. DLS measurements of Z-Avg (d.nm) and PDI for prolate polymersomes persisting over 7 days.

Interestingly, this loss of inner volume does not result in a lower degree of encapsulation for hydrophilic or hydrophobic molecules, but rather a higher degree of retainment. Nile Red and Bovine Serum Albumin(BSA)-Fluorescein were used as model hydrophobic and hydrophilic drugs, respectively. In general, Nile red is encapsulated at a lower efficiency than BSA-Fluorescein due to the smaller hydrophobic volume available in the membrane in comparison to the hydrophilic core. It is assumed that prior to dialysis and shape change, all samples contain the same internal concentration of encapsulated molecules; therefore, the final internal concentration for both molecules must be a result of the polymersomes’ ability to retain the molecules (Table 2). Therefore, we believe the loss of inner volume is not what is responsible for the higher concentration of BSA-Fluorescein and Nile Red in prolate samples, but rather that the NaCl plays some role in the retaining of encapsulated molecules, based on studies of BSA encapsulation in microspheres^40^ and studies that show salts are capable of complexing with payloads to prevent leakage in liposomes^41^. However, this was not investigated further in our system, but might be an interesting subject for future studies.

**Table 2.**
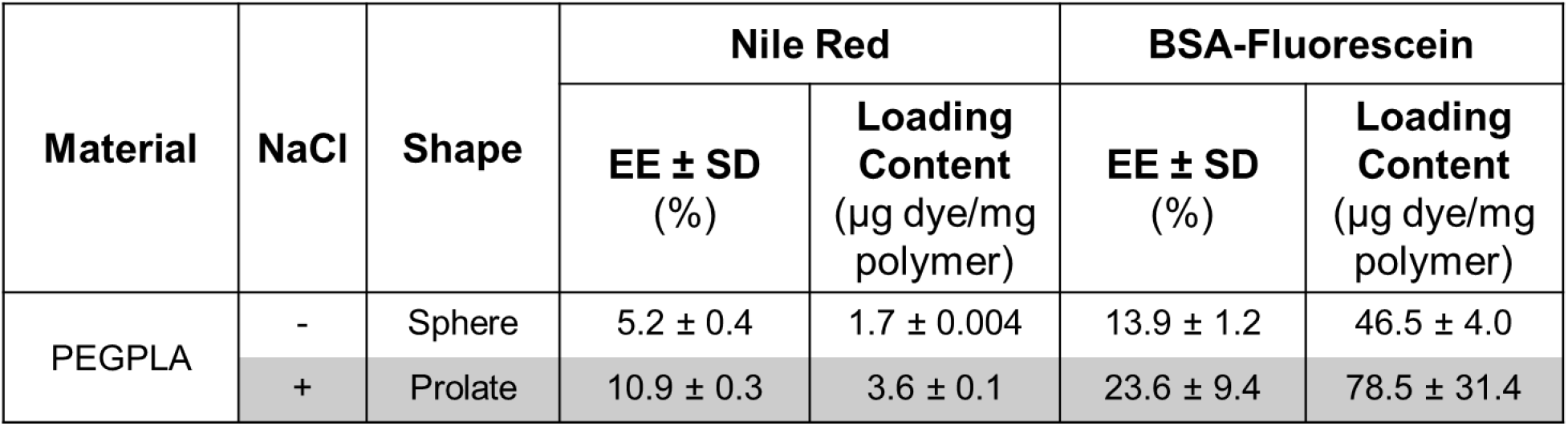
Polymersomes Encapsulation of Multiple Compounds. Retainment of encapsulated dye (Nile Red and Albumin-Fluorescein) by PEGPLA nanoparticles following dialysis against MilliQ Water and NaCl for Spheres and Prolates, respectively.

Prolate shaped particles appear to deliver both BSA-Fluorescein and Nile Red more effectively to SH-SY5Y neuroblastoma cells based on fluorescent images (Figure 6). Intensity of both Nile Red and BSA-Fluorescein is greater when prolate polymersomes were incubated with SH-SY5Y cells. This may be due in part to the higher retainment of both compounds (Table 2). However, this may result from a tendency for cells to take up non-spherical nanoparticles more effectively than spherical nanoparticles, an attribute of the flattened prolate shape granting a greater surface area to contact the cellular surface. Analysis of SH-SY5Y uptake of BSA-Fluorescein by flow cytometry (Figure 7) shows significantly more dye was taken up by SH-SY5Y cells when encapsulated in polymersomes compared to free dye, regardless of shape. The difference in dye uptake between polymersome shapes is trending toward significance, with SH-SY5Y cells appearing to have slightly more internal fluorescence when treated with prolate polymersomes. To make these results more conclusive, more cells could be seeded for analysis as SH-SY5Y cells are partially adherent. SH-SY5Y cells are frequently differentiated into neurons for uptake studies^42-45^, which may also increase adherence. Uptake differences may be due to the change in aspect ratio from spheres to prolates is not large enough to have a significant effect on cell uptake. As demonstrated by Huang, et al, the larger the aspect ratio, the greater than internalization rate with regards to silica-based systems.^46^ To confirm this hypothesis, further work needs to be performed on various self-assembled aspect ratios.

**Figure 6:**
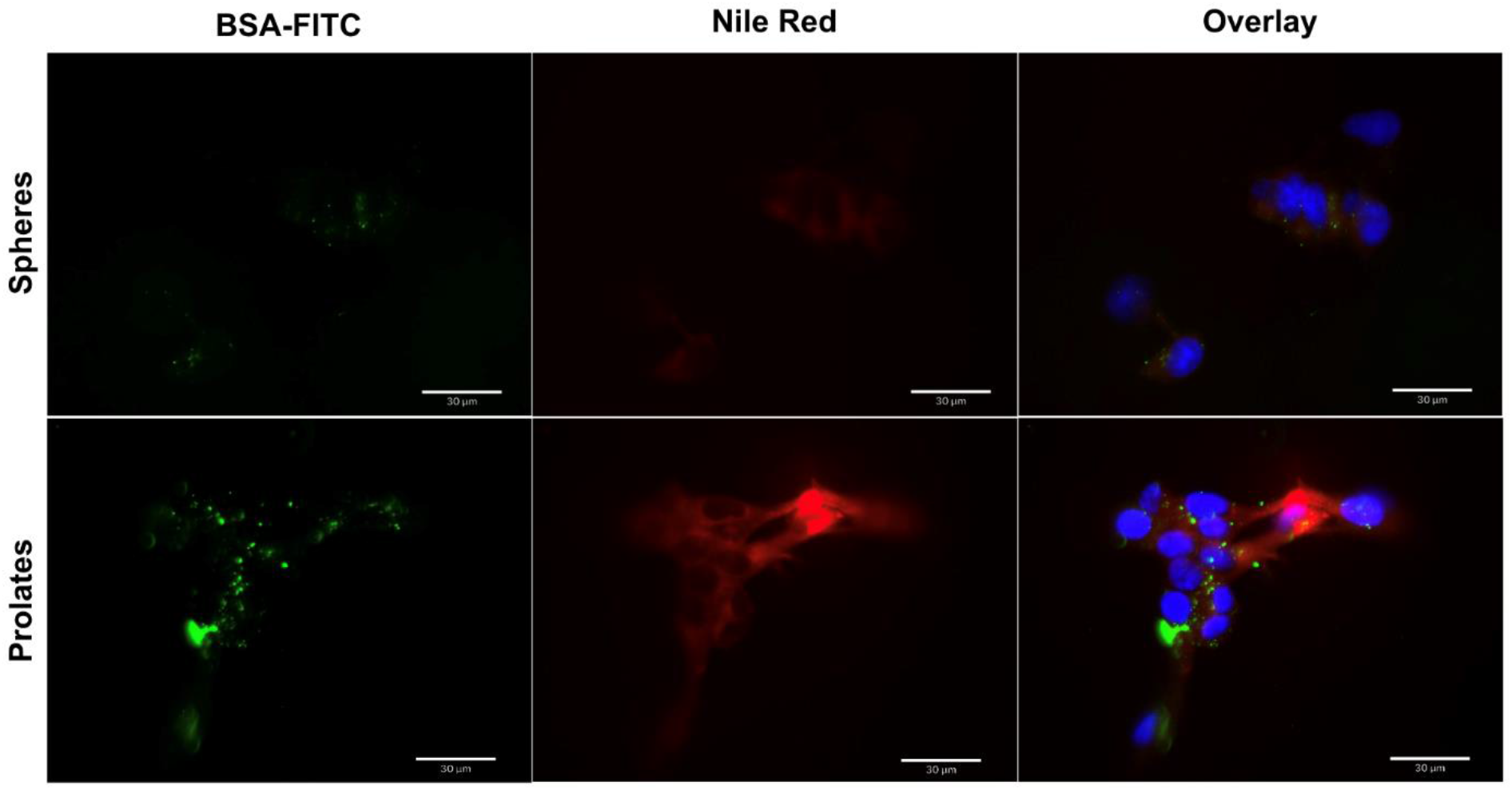
Cell Uptake of Polymersomes Containing Multiple Compounds. Fluorescent images of SH-SY5Yuptake of BSA-Fluorescein (green) and Nile Red (red) after incubation with either spherical or prolate polymersomes for four hours. Nucleus stained with DAPI (blue).

**Figure 7:**
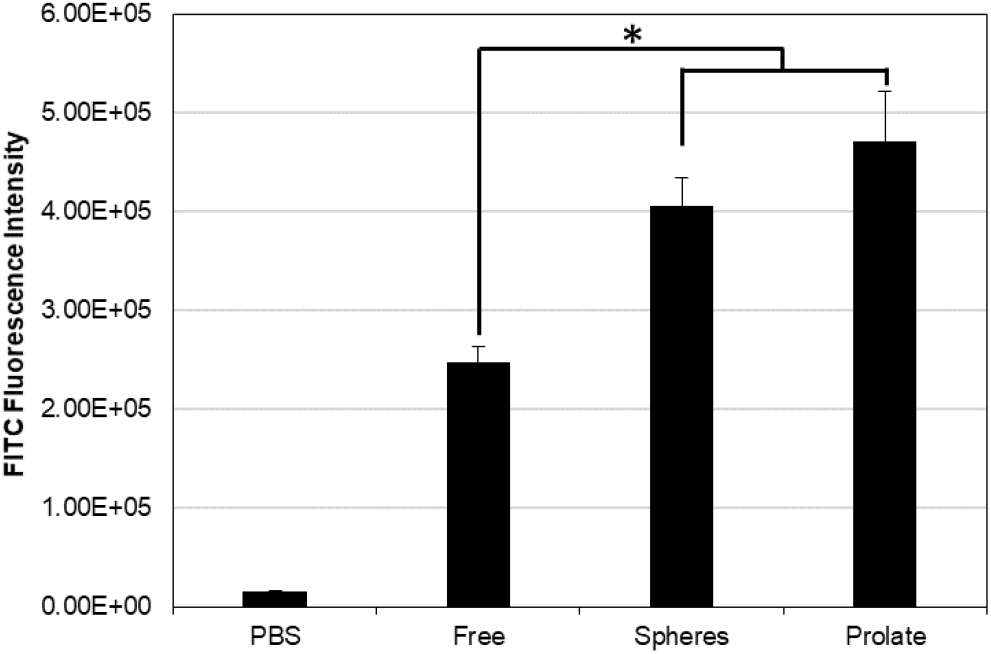
Cell Uptake of Polymersomes Containing Multiple Compoundsconfirmed with Flow Cytometry. After 72 hours, SH-SY5Y cells were analyzed in flow cytometry. Both sphericaland prolate polymersomes lead to greater uptake than free dye (p<0.05). Prolates are trending towards increased uptake compared to spherical polymersomes.

In summary, PEGPLA polymersomes are able to be shifted from a spherical to a more prolate-like shape. To our knowledge, this is the first time self-assembled prolates have been compared to self-assembled spheres with regards to simultaneous hydrophobic and hydrophilic drug delivery. Prolates are capable of encapsulating more of both BSA-fluorescein and Nile Red than their spherical counterparts, which could contribute to the increased intracellular fluorescence observed when cultured with SH-SY5Y neuroblastoma cells. Increasing aspect ratios may be able to increase uptake efficiencies, based on the only slight difference in uptake seen in flow cytometry. Through modulating nanoparticle shape, we are able to enhance uptake of both hydrophilic and hydrophobic drugs, which provides promising results towards efficiently delivering payloads through the blood-brain barrier.

## METHODS

### Polymersome Preparation

Monodisperse spherical polymersomes of Polyethylene glycol (PEG)_1000Da_-b-Polylactic acid (PLA)_5000Da_ (Polyscience, Inc.) were obtained by dissolving the polymer in dimethyl sulfoxide (DMSO) at 15mg/mL then infusing 100μl of the polymer solution into 1mL of MilliQ water at 5μl/min and stirring at 100rpm using a Legato Syringe Pump. For spherical samples, polymersomes were dialyzed in 300 kDa membrane from Spectrum Laboratories against MilliQ water for 15 hours to remove DMSO. Polymersome size and polydispersity were characterized by dynamic light scattering using Malvern Zetasizer Nano ZS.

### Shape Modulation

Upon assembly, polymersomes are spherical and in isotonic conditions. To achieve prolate morphology, samples were subjected to hypertonic conditions. Following infusion, samples, samples were dialyzed against 50mM NaCl solution for 15 hours. To minimize interference, all TEM and SEM imaging was done after an additional dialysis of prolate samples against MilliQ water to remove the salt. TEM and SEM imaging was done with HT7830 Transmission Electron Microscope and High-resolution Scanning Electron Microscope S4800, respectively.

### Encapsulation

Polymersomes loaded with hydrophobic Nile Red were obtained by incorporating Nile Red in the polymer solution prior to infusion at a ratio of 100μg Nile Red per 1mg of PEGPLA. Encapsulation of hydrophilic BSA-Fluorescein was achieved by infusing the polymer solution into a 1mL aqueous solution of BSA-Fluorescein at a concentration of 1mg/mL. Following encapsulation, free dye was removed from samples by 300 kDa membrane dialysis tubes in either MilliQ water or 50mM NaCl, for spheres or prolates, respectively. Following dialysis, samples were removed from dialysis tubing and transferred to an Amicon 100kDa microcentrifuge filter unit and centrifuged in Eppendorf Centrifuge 5424 R for 15 minutes at 5000 RCF. To quantify encapsulated dye, filters were flipped and centrifuged into 200μl DMSO to dissolve nanoparticles and release encapsulated dye. Fluorescent intensity of the dissolved sample was read for BSA-Fluorescein (486nm excitation – 525nm emission) and Nile Red (550nm excitation – 640nm emission) with BioTek Synergy H1M in black clear bottom 96 well assay plates from Corning. Nile Red and BSA-Fluorescein were purchased from Sigma-Aldrich.

### SH-SY5Y Culture and Administration

SH-SY5Y Neuroblastoma cells were cultured in 250ml Falcon flasks at 37 degrees Celsius and 5% carbon dioxide. The media used was a 1:1 mixture of Ham’s F12 and EMEM along with 2mM L-Glutamine, 1% Non-Essential Amino Acid, and 15% Fetal Bovine Serum. The cells were given one week between passage along with two media changes in between.

### SH-SY5Y Fluorescent Imaging

SH-SY5Y cells were added to chamber slides with a seeding density of 93,500 cells per well. The slides were allowed to grow for 72 hours then washed with PBS. Free-dye solutions were prepared at a concentration of 5.4 μg/ml Nile Red and 112.5 μg/ml BSA-Fluorescein, based on encapsulation data (Table 2). In order to prepare the slides for imaging, free-dye solution, suspension of loaded spherical polymersomes and suspension of loaded prolate polymersomes were administered to wells at 25% of the working volume for four hours. After four hours, the slides were prepared for fluorescent imaging. Vectashield mounting medium with DAPI was added to slides prior to drying overnight. Fluorescent images were taken with Echo Revolve hybrid microscope.

### Flow Cytometry with Cytoflex

Prepared 24-well plate with 50,000 cells per well and grown for 48 hours to 70 % confluence. The media was removed and washed out with PBS. We administered 450μl of media and 50μl of PBS for negative control, 450μl of media and 50μl of free-dye solution for positive control, 450μl of media and 50μl of spherical nanoparticles loaded with BSA-Fluorescein, and 450μl of media and 50μl of prolate nanoparticles loaded with BSA-Fluorescein. After 4 hours, the cells were washed three times with cold PBS. They were then trypsinized for one minute with 100μl of 25% trypsin then removed and replaced with 400μl of PBS. Each sample was analyzed with Beckman Coulter CytoFlex Flow Cytometer for 10,000 events.

### Statistical Methods

All statistics were performed using a two-tailed t-test with unequal variance. Statistical significance was defined as p<0.05.

## ACKNOWLEDGEMENTS

We would like to acknowledge Dr. James Morris and Christina Wilkinson for assistance with flow cytometry (COBRE Grant P20GM109094). This work was supported in part by the National Science Foundation EPSCoR Program under NSF Award # OIA-1655740. Any Opinions, findings and conclusions or recommendations expressed in this material are those of the author(s) and do not necessarily reflect those of the National Science Foundation. This work was also partially supported by Clemson’s Creative Inquiry Program.

## DATA AVAILABILITY

The raw/processed data required to reproduce these findings cannot be shared at this time as the data also forms part of an ongoing study.

## REFERENCES

(1) Larsen, J. M.; Martin, D. R.; Byrne, M. E. Recent Advances in Delivery through the Blood-Brain Barrier. Curr. Top. Med. Chem. 2014, No. 14, 1148–1160.

(2) Pardridge, W. M. Drug Delivery to the Brain. J. Cereb. Blood Flow Metab. 1997, No. 17, 713–731.

(3) Huynh, G. H.; Deen, D. F.; Szoka, F. C. Barriers to Carrier Mediated Drug and Gene Delivery to Brain Tumors. J. Control. Release 2006, 110 (2), 236–259. https://doi.org/10.1016/j.jconrel.2005.09.053.

(4) Tsuji, A.; Tamai, I. Blood-Brain Barrier Transport of Drugs. In Introduction to the Blood-Brain Barrier: Methodology, Biology, and Pathology2; Pardridge, W. M., Ed.; Cambridge University Press, 2006; pp 238–245.

(5) Pardridge, W. M. Blood-Brain Barrier Delivery. Drug Discov. Today 2007, 12 (1–2), 54–61. https://doi.org/10.1016/j.drudis.2006.10.013.

(6) Stenehjem, D. D.; Hartz, A. M. S.; Bauer, B.; Anderson, G. W. Novel and Emerging Strategies in Drug Delivery for Overcoming the Blood-Brain Barrier. Future Med. Chem. 2009, 1 (9), 1623–1641. https://doi.org/10.4155/fmc.09.137.

(7) Nathanson, D.; Mischel, P. S. Charting the Course across the Blood-Brain Barrier. J. Clin. Invest. 2011, 121 (1), 31–33. https://doi.org/10.1172/JCI45758.

(8) Pardridge, W. M. Drug Transport across the Blood-Brain Barrier. J. Cereb. Blood Flow Metab. 2012, 32 (11), 1959–1972. https://doi.org/10.1038/jcbfm.2012.126.

(9) Krol, S. Challenges in Drug Delivery to the Brain: Nature Is against Us. J. Control. Release 2012, 164 (2), 145–155. https://doi.org/10.1016/j.jconrel.2012.04.044.

(10) Abbott, N. J. Blood-Brain Barrier Structure and Function and the Challenges for CNS Drug Delivery. J. Inherit. Metab. Dis. 2013, 36 (3), 437–449. https://doi.org/10.1007/s10545-013-9608-0.

(11) Bell, R. D.; Ehlers, M. D. Breaching the Blood-Brain Barrier for Drug Delivery. Neuron 2014, 81 (1), 1–3. https://doi.org/10.1016/j.neuron.2013.12.023.

(12) Jain, R. K.; Janda, K. D.; Saltzman, W. M. Drug Discovery and Delivery. Mol. Med. Today 1995, 1 (1), 4. https://doi.org/10.1016/1357-4310(95)80005-0.

(13) Discher, D. E.; Ahmed, F. Polymersomes. Annu. Rev. Biomed. Eng. 2006, 8, 323–341. https://doi.org/10.1146/annurev.bioeng.8.061505.095838.

(14) Onaca, O.; Enea, R.; Hughes, D. W.; Meier, W. Stimuli-Responsive Polymersomes as Nanocarriers for Drug and Gene Delivery. Macromol. Biosci. 2009, 9 (2), 129–139. https://doi.org/10.1002/mabi.200800248.

(15) Ahmed, F.; Pakunlu, R. I.; Srinivas, G.; Brannan, A.; Bates, F.; Klein, M. L.; Minko, T.; Discher, D. E. Shrinkage of a Rapidly Growing Tumor by Drug-Loaded Polymersomes: PH-Triggered Release through Copolymer Degradation. Mol. Pharm. 2006, 3 (3), 340–350. https://doi.org/10.1021/mp050103u.

(16) Popescu, M.; Tsitsilianis, C. Controlled Delivery of Functionalized Gold Nanoparticles by PH-Sensitive Polymersomes. Support. Inf. 5000, 0–7.

(17) Meng, F.; Zhong, Z.; Feijen, J. Stimuli-Responsive Polymersomes for Programmed Drug Delivery. Biomacromolecules 2009, 10 (2), 197–209. https://doi.org/10.1021/bm801127d.

(18) Ahmed, F.; Pakunlu, R. I.; Brannan, A.; Bates, F.; Minko, T.; Discher, D. E. Biodegradable Polymersomes Loaded with Both Paclitaxel and Doxorubicin Permeate and Shrink Tumors, Inducing Apoptosis in Proportion to Accumulated Drug. J. Control. Release 2006, 116 (2), 150–158. https://doi.org/10.1016/j.jconrel.2006.07.012.

(19) Jain, J. P.; Kumar, N. Self Assembly of Amphiphilic (PEG)(3)-PLA Copolymer as Polymersomes: Preparation, Characterization, and Their Evaluation as Drug Carrier. Biomacromolecules 2010, 11 (4), 1027–1035. https://doi.org/10.1021/bm1000026.

(20) Kim, J.-K.; Garripelli, V. K.; Jeong, U.-H.; Park, J.-S.; Repka, M. A.; Jo, S. Novel PH-Sensitive Polyacetal-Based Block Copolymers for Controlled Drug Delivery. Int. J. Pharm. 2010, 401 (1–2), 79–86. https://doi.org/10.1016/j.ijpharm.2010.08.029.

(21) Kelly, J. M.; Gross, A. L.; Martin, D. R.; Byrne, M. E. Polyethylene Glycol-b-Poly(Lactic Acid) Polymersomes as Vehicles for Enzyme Replacement Therapy. Nanomedicine 2017, 12 (23), 2591–2606. https://doi.org/10.2217/nnm-2017-0221.

(22) Yu, J.; Qian, C.; Zhang, Y.; Cui, Z.; Zhu, Y.; Shen, Q.; Ligler, F. S.; Buse, J. B.; Gu, Z. Hypoxia and H 2 O 2 Dual-Sensitive Vesicles for Enhanced Glucose-Responsive Insulin Delivery. Nano Lett. 2017, acs.nanolett.6b03848. https://doi.org/10.1021/acs.nanolett.6b03848.

(23) Dai, L.; Cai, R.; Li, M.; Luo, Z.; Yu, Y.; Chen, W.; Shen, X.; Pei, Y.; Zhao, X.; Cai, K. Dual-Targeted Cascade-Responsive Prodrug Micelle System for Tumor Therapy in Vivo. Chem. Mater. 2017, acs.chemmater.7b02513. https://doi.org/10.1021/acs.chemmater.7b02513.

(24) Hu, X.; Yu, J.; Qian, C.; Lu, Y.; Kahkoska, A. R.; Xie, Z.; Jing, X.; Buse, J. B.; Gu, Z. H2O2-Responsive Vesicles Integrated with Transcutaneous Patches for Glucose-Mediated Insulin Delivery. ACS Nano 2017, 11 (1), 613–620. https://doi.org/10.1021/acsnano.6b06892.

(25) Kulkami, P.; Mallik, S. Drug Delivery Vehicles from Stimuli-Responsive Block Copolymers. In Novel Nanoscale Hybrid Materials; Chauhan, B. P. S., Ed.; John Wiley & Sons, 2018; pp 239–261.

(26) Banerjee, A.; Qi, J.; Gogoi, R.; Wong, J.; Mitragotri, S. Role of Nanoparticle Size, Shape and Surface Chemistry in Oral Drug Delivery. 2016, 238 (805), 176–185. https://doi.org/10.1016/j.jconrel.2016.07.051.

(27) Hao, N.; Li, L.; Zhang, Q.; Huang, X.; Meng, X.; Zhang, Y.; Chen, D.; Tang, F.; Li, L. The Shape Effect of PEGylated Mesoporous Silica Nanoparticles on Cellular Uptake Pathway in Hela Cells. Microporous Mesoporous Mater. 2012, 162, 14–23. https://doi.org/10.1016/j.micromeso.2012.05.040.

(28) Jo, D. H.; Kim, J. H.; Lee, T. G.; Kim, J. H. Size, Surface Charge, and Shape Determine Therapeutic Effects of Nanoparticles on Brain and Retinal Diseases. Nanomedicine Nanotechnology, Biol. Med. 2015, 11 (7), 1603–1611. https://doi.org/10.1016/j.nano.2015.04.015.

(29) Kolhar, P.; Anselmo, A. C.; Gupta, V.; Pant, K.; Prabhakarpandian, B.; Ruoslahti, E.; Mitragotri, S. Using Shape Effects to Target Antibody-Coated Nanoparticles to Lung and Brain Endothelium. Proc. Natl. Acad. Sci. 2013, 110 (26), 10753–10758. https://doi.org/10.1073/pnas.1308345110.

(30) Abdelmohsen, L. K. E. A., Williams, D. S., Pille, J., Ozel, S. G., Rikken, R. S. M., Wilson, D. A. & van Hest, J. C. M. Formation of Well-Defined, Functional Nanotubes via Osmotically Induced Shape Transformation of Biodegradable Polymersomes. J. Am. Chem. Soc. 138, 9353–9356 (2016).

(31) Rikken, R. S. M., Engelkamp, H., Nolte, R. J. M., Maan, J. C., Van Hest, J. C. M., Wilson, D. A. & Christianen, P. C. M. Shaping polymersomes into predictable morphologies via out-of-equilibrium self-assembly. Nat. Commun. 7, 1–7 (2016).

(32) Salva, R., Le Meins, J. F., Sandre, O., Bruîlet, A., Schmutz, M., Guenoun, P. & Lecommandoux, S. Polymersome shape transformation at the nanoscale. ACS Nano 7, 9298–9311 (2013).

(33) Men, Y., Li, W., Janssen, G.-J. A., Rikken, R. S. M. & Wilson, D. A. “Stomatocyte in stomatocyte: a new shape of polymersome induced via chemical addition methodology”. Nano Lett. acs.nanolett.8b00187 (2018). doi:10.1021/acs.nanolett.8b00187

(34) Wauters, A. C., Pijpers, I. A. B., Mason, A. F., Williams, D. S., Tel, J., Abdelmohsen, L. K. E. A. & Van Hest, J. C. M. Development of Morphologically Discrete PEG-PDLLA Nanotubes for Precision Nanomedicine. 20, 16 (2019).

(35) Abdelmohsen, L. K. E. A., Rikken, R. S. M., Christianen, P. C. M., van Hest, J. C. M. & Wilson, D. A. Shape characterization of polymersome morphologies via light scattering techniques. Polymer (Guildf). 107, 445–449 (2016).

(36) Bhattacharjee, S. DLS and zeta potential - What they are and what they are not? J.Control. Release 235, 337–351 (2016).

(37) Sánchez-Purrà, M.; Ramos, V.; Petrenko, V. A.; Torchilin, V. P.; Borrós, S. Double-Targeted Polymersomes and Liposomes for Multiple Barrier Crossing. Int. J. Pharm. 2016, 511 (2), 946–956. https://doi.org/10.1016/j.ijpharm.2016.08.001.

(38) Trickler, W. J.; Lantz, S. M.; Murdock, R. C.; Schrand, A. M.; Robinson, B. L.; Newport, G. D.; Schlager, J. J.; Oldenburg, S. J.; Paule, M. G.; Slikker, W.; et al. Silver Nanoparticle Induced Blood-Brain Barrier Inflammation and Increased Permeability in Primary Rat Brain Microvessel Endothelial Cells. Toxicol. Sci. 2010, 118 (1), 160–170. https://doi.org/10.1093/toxsci/kfq244.

(39) Ivask, A.; Pilkington, E. H.; Blin, T.; Kakinen, A.; Vija, H.; Visnapuu, M.; Quinn, J. F.; Whittaker, M. R.; Qiao, R.; Davis, T. P.; et al. Uptake and Transcytosis of Functionalized Superparamagnetic Iron Oxide Nanoparticles in an in Vitro Blood Brain Barrier Model. Biomater. Sci. 2017. https://doi.org/10.1039/C7BM01012E.

(40) Pistel, K. F.; Kissel, T. Effects of Salt Addition on the Microencapsulation of Proteins Using W/O/W Double Emulsion Technique. J. Microencapsul. 2000, 17 (4), 467–483.

(41) Gubernator, J.; Chwastek, G.; Korycińska, M.; Stasiuk, M.; Grynkiewicz, G.; Lewrick, F.; Süss, R.; Kozubek, A. The Encapsulation of Idarubicin within Liposomes Using the Novel EDTA Ion Gradient Method Ensures Improved Drug Retention in Vitro and in Vivo. J. Control. Release. 2010, 146 (1), 68–75. https://doi.org/10.1016/j.jconrel.2010.05.021.

(42) Encinas, M.; Iglesias, M.; Liu, Y.; Wang, H.; Muhaisen, A.; Ceña, V.; Gallego, C.; Comella, J. X. Sequential Treatment of SH-SY5Y Cells with Retinoic Acid and Brain-Derived Neurotrophic Factor Gives Rise to Fully Differentiated, Neurotrophic Factor-Dependent, Human Neuron-Like Cells. J. Neurochem. 2002, 75 (3), 991–1003. https://doi.org/10.1046/j.1471-4159.2000.0750991.x.

(43) Shipley, M. M.; Mangold, C. A.; Szpara, M. L. Differentiation of the SH-SY5Y Human Neuroblastoma Cell Line. JoVE. 2016, No. 108, e53193. https://doi.org/doi:10.3791/53193.

(44) Lopes, F. M.; Schröder, R.; Júnior, M. L. C. da F.; Zanotto-Filho, A.; Müller, C. B.; Pires, A. S.; Meurer, R. T.; Colpo, G. D.; Gelain, D. P.; Kapczinski, F.; et al. Comparison between Proliferative and Neuron-like SH-SY5Y Cells as an in Vitro Model for Parkinson Disease Studies. Brain Res. 2010, 1337 (February 2018), 85–94. https://doi.org/10.1016/j.brainres.2010.03.102.

(45) Schneider, L.; Giordano, S.; Zelickson, B. R.; S Johnson, M.; A Benavides, G.; Ouyang, X.; Fineberg, N.; Darley-Usmar, V. M.; Zhang, J. Differentiation of SH-SY5Y Cells to a Neuronal Phenotype Changes Cellular Bioenergetics and the Response to Oxidative Stress. Free Radic. Biol. Med. 2011, 51 (11), 2007–2017. https://doi.org/10.1016/j.freeradbiomed.2011.08.030.

(46) Huang, X.; Teng, X.; Chen, D.; Tang, F.; He, J. The Effect of the Shape of Mesoporous Silica Nanoparticles on Cellular Uptake and Cell Function. Biomaterials. 2010, 31 (3), 438–448. https://doi.org/10.1016/j.biomaterials.2009.09.060.

